# High-resolution mapping of *Ryd4^Hb^*, a major resistance gene to *Barley yellow dwarf virus* from *Hordeum bulbosum*

**DOI:** 10.1101/2023.09.19.558385

**Authors:** Hélène Pidon, Brigitte Ruge-Wehling, Torsten Will, Antje Habekuß, Neele Wendler, Klaus Oldach, Anja Maasberg-Prelle, Viktor Korzun, Nils Stein

## Abstract

Virus diseases are causing high yield losses in crops worldwide. The *Barley yellow dwarf virus* (BYDV) complex is responsible for one of the most widespread and economically important viral diseases of cereals. While no complete resistance gene has been uncovered in the primary genepool of barley, sources of resistance were identified in the wild relative *Hordeum bulbosum*, representing the secondary genepool of barley. One such locus, *Ryd4*^*Hb*^, has been previously introgressed into barley, and was allocated to chromosome 3H, but is tightly linked to a sublethality factor that prevents the incorporation and utilization of *Ryd4*^*Hb*^ in barley varieties. To solve this problem, we fine-mapped *Ryd4*^*Hb*^ and separated it from this negative factor. We narrowed the *Ryd4*^*Hb*^ locus to a 66.5 kbp physical interval in the barley ‘Morex’ reference genome. The region comprises one complete and one partial gene from the nucleotide-binding and leucine-rich repeat immune receptor family, typical of dominant virus resistance genes. The closest homolog to these two *Ryd4*^*Hb*^ candidate genes is the wheat *Sr35* stem rust resistance gene. In addition to the fine mapping, we reduced the sublethality factor interval to 600 kbp in barley. Aphid feeding experiments demonstrated that *Ryd4*^*Hb*^ provides a direct resistance to BYDV rather than a resistance to its vector. The presented results, including the high-throughput molecular markers, will permit a more targeted selection of the resistance in breeding, enabling the use of *Ryd4*^*Hb*^ in barley varieties.

**Key message:** We mapped *Ryd4*^*Hb*^ in a 66.5 kpb interval in barley and dissociated it from a sublethality factor. These results will enable a targeted selection of the resistance in barley breeding.

## Introduction

Virus diseases cause significant yield losses and represent an increasing threat to agricultural crop production worldwide (Oerke, 2006). Among them, *Barley yellow dwarf virus* (BYDV) complex is responsible for one of the most widespread and economically important viral diseases of cereals. Transmitted in a persistent and circulative manner by several species of aphids, BYDV causes dwarfing and leaf discoloration, leading to significant yield loss in major cereal crops, in particular barley, wheat, maize, and oats (Ali et al., 2018). In recent years, it has become increasingly important in winter barley with an incidence that could reach 70% and yield loss of up to 80% (Beoni et al., 2016; Dedryver et al., 2010; Ordon et al., 2009). As climate change scenarios predict longer and warmer autumns, which favor aphid infestations of winter crop fields, BYDV could become one of the most threatening diseases of cereal crops (Roos et al., 2011; Trebicki, 2020). Reduction of yield losses by insecticide-based vector control is possible in principle, but undesirable for ecological reasons. To ensure sustainable barley cultivation in the expanding infestation areas and thus secure yields and quality in the long term, the cultivation of virus resistant varieties would provide the best solution.

So far, three genes and some QTLs have been described as providing partial resistance or tolerance to BYDV in barley. The gene *ryd1*, providing recessive intermediate tolerance was identified by Suneson (1955) but is still not cloned (Niks et al., 2004). Its effectiveness is low and it is rarely used in breeding. *Ryd2* was identified from an Ethiopian barley landrace (Schaller et al., 1964). It provides field tolerance to the virus serotypes BYDV-PAV, BYDV-MAV, and BYDV-SGV (Baltenberger et al., 1987). Mapped close to the centromere of chromosome 3H (Collins et al., 1996), *Ryd2* is used in several breeding lines (Kosova et al., 2008) where it can reduce significantly the yield loss caused by BYDV (Beoni et al., 2016). The third gene, *Ryd3* was also identified from an Ethiopian barley landrace (Niks et al., 2004). The gene was mapped in the centromeric region of chromosome 6H but, despite fine mapping on more than 3,000 F_2_ plants (Lüpken et al., 2014), the mapping interval is still large. *Ryd3* has been transferred to commercial varieties where it provides a quantitative resistance, improved when in combination with *Ryd2* (Riedel et al., 2011). QTLs on chromosomes 1H, 2H, 4H, 5H, and 7H have been reported, however, providing only a limited level of tolerance (Toojinda et al., 2000; Riedel et al., 2011; Hu et al., 2019). No complete resistance to BYDV or its aphid vectors is known in barley, and broadening the genetic basis of resistance is therefore needed to ensure a durable and stable production of winter barley fields.

The secondary genepool of barley, consisting of the species *Hordeum bulbosum*, has not yet been used to improve resistance to the BYDV complex. Michel (1996) identified resistance to BYDV in the tetraploid (2n=4x=28) *Hordeum bulbosum* accession A17 (Bu10/2) from the Botanical Garden of Montevideo, Uruguay. Plants of this accession remained ELISA-negative for BYDV after several inoculations with aphids charged with the virus isolates BYDV-PAV1 Aschersleben, BYDV-MAV1 Aschersleben, and CYDV (*Cereal yellow dwarf virus*)-RPV Dittersbach (Habekuß et al., 2004). A17 was used as a parent in interspecific crosses and backcrosses with *H. vulgare* cv. Igri to generate an *H. bulbosum* introgression to barley. Its resistance was described as complete, dominant, and monogenic, and the locus, assigned to chromosome 3H, was named *Ryd4*^*Hb*^ (Scholz et al., 2009). Adversely, a recessive sublethality factor was cosegregating with *Ryd4*^*Hb*^ in the respective introgression. A study revealed low aphid feeding on the *H. bulbosum* A17 accession, suggesting that resistance may not be acting directly on the virus but rather indirectly against the aphid vector (Schliephake et al., 2013).

The present study reports the fine mapping of the *Ryd4*^*Hb*^ locus, the identification of candidate genes, and the description of aphids feeding behavior on susceptible and resistant introgression lines.

## Material and methods

### Plant material

BC_2_F_5_ and BC_2_F_6_ selected from BC_2_F_4_ plants from the Scholz et al. (2009) population were used for the low-resolution linkage mapping and development of an introgression line lacking the sublethality factor. This population is later named LM_Pop.

Two additional populations were generated to map *Ryd4*^*Hb*^ at a higher resolution. FM_Pop1 was derived from a BC_2_F_7_ plant from LM_Pop crossed successively with three different barley elite varieties. The pedigree of the 15 lineages that constitute FM_Pop1 is presented in supplementary table 1. The donor of resistance in the FM_Pop2 was a BC_2_F_8_ derived from the BC_2_F_6_ JKI-5215 homozygous for the *H. vulgare* (Hv)-allele in the sublethality factor locus. As for FM_Pop1, it was crossed successively with two different barley elite varieties. The pedigree of the lines of four lineages that constitute FM_Pop2 is presented in supplementary table 2. The F_1_ plants from the successive crosses were checked with markers to ensure the presence of the *H. bulbosum* (Hb)-allele at the *Ryd4*^*Hb*^ locus and were further selfed to obtain the F_2_ lineages forming FM_Pop1 and FM_Pop2.

**Table 1.**
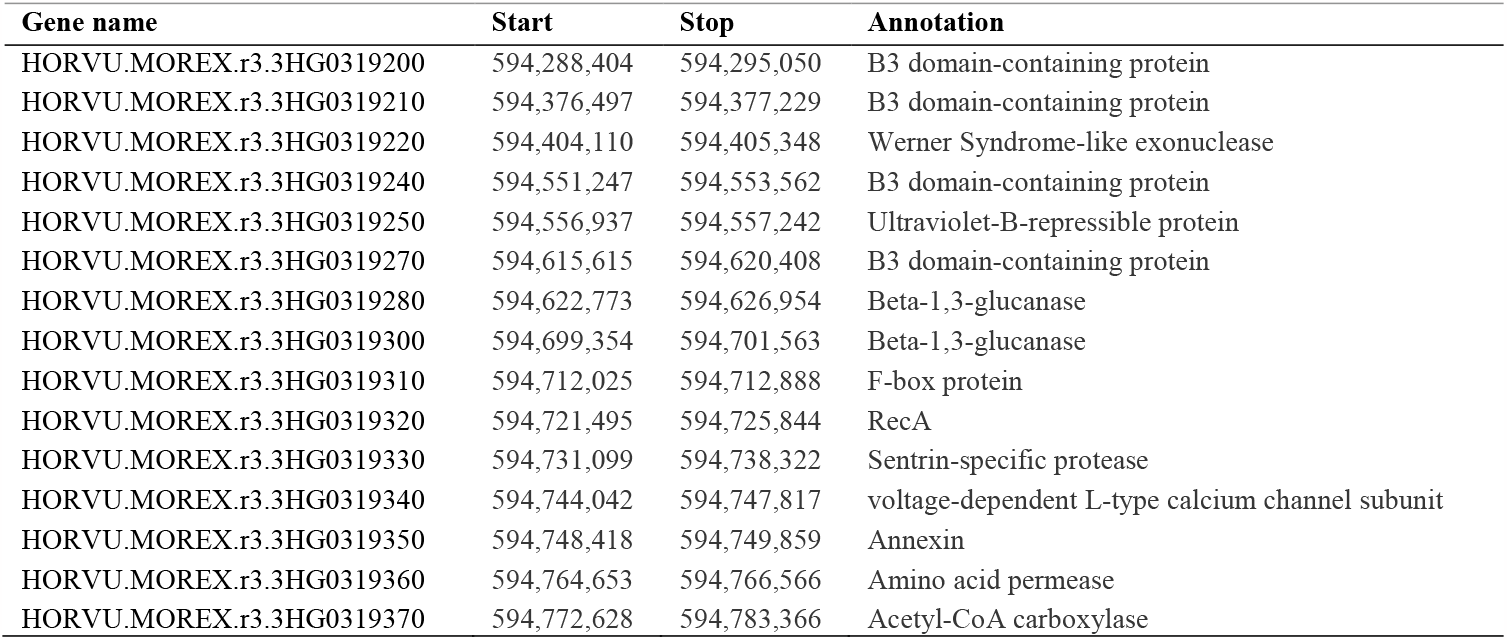
High-confidence genes annotated in the sublethality interval on MorexV3 reference genome. Coordinates refer to chromosome 3H.

### Test of resistance to BYDV

Five to ten *Rhopalosiphum padi* aphids of the biotype R07, that were reared on BYDV-PAV1 Aschersleben (PAV1-ASL) infected plants as described by Kern et al. (2022), were placed on each one-week-old seedlings to be phenotyped in an air-conditioned greenhouse (20 °C, 16 h photoperiod, 10 klx). The aphids were killed after two days using the insecticide Confidor ® WG 70 (Bayer CropScience AG, Germany). Further cultivation of the plants was carried out. Five to six weeks after inoculation, leaf samples of 50 mg from two leaves were taken and tested by a double-antibody sandwich-enzyme-linked immunosorbent assay (DAS-ELISA) according to Clark & Adams (1977) using custom-made antibodies. The viral content was evaluated by measuring extinction at 405 nm on a microtitre plate reader (Opsys MR, ThermoLabsystems or Tecan Sunrise, Tecan) one hour after the addition of the enzyme substrate. Based on negative controls, a extinction threshold was set in each experiment, usually at 0.1, under which a plant was classified as resistant. Phenotyping tests for gene mapping were performed on ten to 15 seeds of progenies of each genotype.

### DNA extraction

Genomic DNA from the LM-Pop was either isolated following a slightly modified protocol after Stein et al. (2001) or using the Biosprint 96 DNA Plant Kit (Qiagen) and the Biosprint 96 working station (Qiagen) following the manufacturer’s instructions. DNA was dissolved in TE buffer, quantified via photometric approaches (NanoQuant, Tecan, Austria) and diluted to a working concentration of 10 ng/µl. DNA extractions of plants from FM_Pop1 and FM_Pop2 were carried out according to the protocol described in Milner et al. (2019).

### Marker development for low-resolution linkage mapping

The EST-derived SSR anchor markers *GBM1050* and *GBM1059* (Scholz et al., 2009; Stein et al., 2007; Thiel et al., 2003; Varshney et al., 2007) were kindly provided by Prof. Andreas Graner (Leibniz Institute of Plant Genetics and Crop Plant Research, Gatersleben). The STS markers *GLMWG883* and *GLABC161* were derived from the sequence of RFLP probe *MWG883* (Rostoks et al., 2005; Szűcs et al., 2009) and barley anchor marker *ABC161* (Close et al., 2009), respectively. Orthology of the interval on the rice chromosome 1 using the mapping of the anchor marker *ABC161* at 40.43 Mbp on said chromosome allowed for the development of 18 tentative consensus (TC) markers polymorphic between Hv and Hb on chromosome 3HL.

Additional polymorphisms between the *H. bulbosum* and *H. vulgare* genomes for marker development were identified by RNASeq and Massive analysis of cDNA-ends (MACE). To this end, 1,000 plants from BC_2_F_5_ of LM_Pop were screened with the TC marker *TC173485*. Among them, 200 plants homozygous for the *H. bulbosum* allele, considered resistant, and 200 plants homozygous for the *H. vulgare* allele, considered susceptible, were selected. In total, 100 plants of each category were inoculated by aphids carrying the isolate BYDV-PAV1, and an equal amount of plants were infested with control aphids without virus titer in a separate climate chamber. 1h, 4h, 8h, and 24h after inoculation plant material of 25 genotypes per category and treatment was harvested and sent to GenXPro (Frankfurt am Main, Germany) for RNA isolation and sequencing. RNASeq and MACE were performed as described in Santos et al. (2018) and Braun et al. (2019), respectively. In short, the raw data was cleaned off adapter sequences using the software TagDust (Lassmann et al., 2009). All RNAseq datasets were combined to create a reference library. Assembly was performed using the software Trinity (Grabherr et al., 2011). The reads of the individual libraries were hereafter mapped to the reference library and single nucleotide polymorphisms (SNPs) were identified using the software SNVMix (Goya et al., 2010). SNPs between the *H. bulbosum* and *H. vulgare* genomes were identified and sequences 100 bps up- and downstream of each SNP were determined. Annotation of the SNP-containing sequences was done by using the database Swiss-Prot (Boeckmann et al., 2003).

Additional polymorphisms were retrieved from exome capture sequencing, performed according to Wendler et al. (2014) on the *H. bulbosum* parent A17 (Wendler et al., 2015) and the BC_2_F_4_ plant 5194/5, homozygous for the 3HL-*H. bulbosum* introgression and mapped on the first barley genome assembly (International Barley Genome Sequencing Consortium, 2012). Single nucleotide variants between *H. vulgare* and *H. bulbosum* were called with samtools (SNP call score <200). Variants located within 200 bp of the end of a reference sequence contig or supported by less than fivefold sequence read coverage were excluded from further evaluation. The flanking sequences (50–60 bp) of variant positions were used for marker assay development.

The primer design for PCR markers was carried out using Primer 3 (Untergasser et al., 2012). Conversion of SNPs in CAPS markers was done by using SNP2CAPS (Thiel et al., 2004). All markers used for the low-resolution linkage mapping are described in supplementary table 3.

### Marker development for high-resolution linkage mapping

Exome capture data of the *H. bulbosum* A17 parent and of the BC_2_F_4_ plant 5194/5 was remapped onto the barley reference genome MorexV3 (Mascher et al., 2021) together with the exome capture data of 13 barley varieties from (Russell et al., 2016) (cultivars ‘Barke’, ‘Bonus’, ‘Borwina’, ‘Bowman’, ‘Foma’, ‘Gull’, ‘Harrington’, ‘Haruna-Nijo’, ‘Igri’, ‘Kindred’, ‘Morex’, ‘Steptoe’, and ‘Vogelsanger Gold’). Reads mapping and variant calling were performed as described in Milner et al., (2019). The SNP matrix was filtered on the following criteria: heterozygous and homozygous calls had to have a minimum mapping quality score of three and five, respectively, and be supported by a minimum of ten reads. SNP sites were retained if they had less than 20% missing data and less than 20% heterozygous calls. SNPs that were within the *Ryd4*^*Hb*^ 20 Mbp interval defined by the low-resolution linkage mapping, homozygous for one allele in all barley varieties and for the other allele in A17 and the BC_2_F_4_ introgression line, were retained.

For six SNPs, a 100 bp sequence containing the SNP in its center was provided to LGC genomics (Berlin, Germany) for KASPar marker production (Supplementary table 4). Within the sublethality factor interval, ten more SNPs were retrieved and sequences of 100 bp around each one were sent to 3CR Bioscience (Welwyn Garden City, UK) for PACE assay design (Supplementary table 5). Primers were ordered from Metabion (Germany) and mixed according to 3CR Bioscience (Welwyn Garden City, UK) recommendations.

Thirteen CAPS markers (Supplementary table 6) were developed using NEBcutter (Vincze et al., 2003) to identify the cutting enzyme and Primer 3 (Untergasser et al., 2012) to design the PCR primers.

### Genotyping assays

For PCR of SSR, STS, and CAPS markers were carried out in a volume of 10µL containing 50-100ng of DNA, 1X PCR buffer (Qiagen), 0.5 mM of each primer, 0.5 U of Taq DNA polymerase (Qiagen), and 0.2 mM of dNTPs. PCR amplification was carried out with an initial 10 min step at 95°C, followed by a touchdown profile of ten cycles at 95°C for 30 seconds, 60°C for 30 seconds with a 0.5°C reduction per cycle, and 72°C for 1 minute, followed by 35 cycles at 95°C for 30 seconds, 55°C for 30 seconds then 72°C for 1 minute, and a last step of 7 minutes at 72°C. For CAPS markers, a 5 µl aliquot of the PCR product was digested in 10 µL with 1 U of restriction enzyme and 1x of the appropriate digestion buffer at the temperature recommended by the manufacturer. Pre- and post-digestion PCR products were separated on 2.5 % agarose gels followed by ethidium bromide staining or in 10% polyacrylamide gels followed by silver nitrate staining according to Budowle et al. (1991).

Detection of SNPs as genetic markers were performed by high-resolution melt analysis (HRM) by using the Rotor Gene Technology (Qiagen). PCR was carried out in 20 µl volume containing 20 ng template DNA, 1 x buffer (Promega), 2.5 mM MgCl_2_, 0.8 mM dNTP mix, 0.5 µM of each primer, 1 x EvaGreen Dye (Biotium, Inc.) and 0.3 U *Taq* DNA polymerase (Promega). A touchdown PCR protocol was conducted with a temperature gradient from 60–50°C. The melt curve analysis was conducted by ramping from 65 C to 95 C with a 0.1 C decrease per capture.

Genotyping assays with KASPar and PACE markers were carried out as described in Pidon et al. (2020).

### Genetic linkage analysis

LM_Pop was genotyped with 4 EST-derived SSR anchor markers, 18 TC-markers, 6 MACE-SNP, 13 STS-markers derived from MACE (denoted as MACE_b), 9 markers derived from RNA-seq experiment, and 3 markers derived from exome capture. Linkage analysis was performed using the JoinMap® 4.1 software (Van Ooijen, 2006). Genetic maps were displayed and edited in MapChart2.2 (Voorrips, 2002).

### Pangenome comparison

The flanking markers and the *Ryd4*^*Hb*^ interval in the MorexV3 genome were searched on the 19 other assemblies of the barley pangenome (Jayakodi et al., 2020) using BLAST+ (Camacho et al., 2009). The resulting intervals were extracted and reannotated through a combination of alignments of the Morex candidate genes to them, the search for the presence of conserved domains using NCBI *conserved domains* (S. Lu et al., 2019), and NLRs annotation with NLR-Annotator (Steuernagel et al., 2020). The interval structures were compared using Easyfig with blastn (Sullivan et al., 2011).

### Aphids feeding experiment

In order to test if *Ryd4*^*Hb*^ provides resistance to aphids, a resistant and a susceptible progeny of the heterozygous lines at *Ryd4Hb* locus EP_16-271_3_206 and EP_16-460_5_228 were selected. The susceptible progenies EP_16-271_3_206_2 and EP_16-460_5_228_2 were carrying Hv alleles at both Ryd4_CAPS19 and Ryd4_CAPS24, while the resistant lines EP_16-271_3_206_4 and EP_16-460_5_228_6 were recombining in the interval and carrying a Hb allele at Ryd4_CAPS24 or Ryd4_CAPS19, respectively. The feeding behavior of 12 to 16 adult apterouse non-viruliferous *R. padi* (clone R07) of random age was observed on an individual healthy plant of the respective genotypes per aphid by using the electrical penetration graph (EPG) technique (Tjallingii, 1978). Plants for EPG experiments were reared in a greenhouse and were used at a 3-4 leaf stage where aphids were placed on the lower side of the second leaf. Aphids were reared as described before (Kern et al., 2022). Aphids starved for 1 hour before they were placed on the leaf. The recording started when all aphids were prepared. The observation period was set to 8 hours and recording was started after all aphids were placed. For data acquisition, the GIGA-8 EPG amplifier and EPG stylet software (EPG Systems, Wageningen, The Netherlands) were used and data were analyzed with the EPG stylet analysis module. Waveforms were annotated according to Tjallingii (1978) and Tjallingii & Esch (1993). Subsequently, selected parameters were analyzed by using an Excel workbook (Alvarez et al., 2021). Recordings of aphids that fell from the leaf or escaped during the experiment were not used.

## Results

### Low-resolution linkage mapping of *Ryd4*^*Hb*^

From LM_Pop, 1,125 BC_2_F_5_ and BC_2_F_6_ were used for the low-resolution linkage mapping of *Ryd4*^*Hb*^. Phenotyping of progenies revealed 570 resistant, 276 susceptible plants, and 279 plants that died at early stages. This high mortality rate was expected due to the recessively inherited sublethality factor very closely linked to *Ryd4*^*Hb*^. This locus prevents normal plant development, resulting in plant death (Figure 1). Because of the close linkage between the two loci, we considered that all plants that died in the BC_2_F_5_ and BC_2_F_6_ families could be defined as homozygous resistant, while the resistant plants that survived would be heterozygous at *Ryd4*^*Hb*^. The phenotype distribution would indeed fit the expected 1:2:1 ratio of homozygous resistant, heterozygous resistant, and homozygous susceptible genotypes for a dominant monogenic inheritance of *Ryd4*^*Hb*^ resistance (Chi-Square Goodness of Fit Test: χ^2^= 0.216, *p*=0.90), confirming the previous observation for that locus on the BC_2_F_4_ generation (Scholz et al., 2009).

**Fig. 1.**
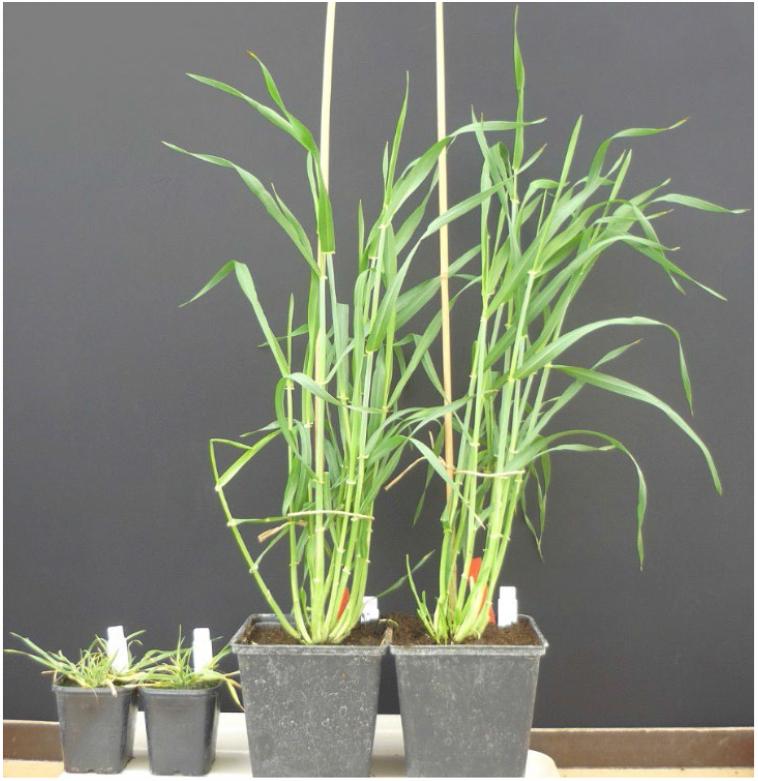
Five months-old homozygous resistant plants either carrying a sublethality factor (left) or vital (right). The growth of plants homozygous at the sublethality factor is greatly reduced compared to plants carrying the resistance locus without the sublethality factor. They died before reaching heading.

The population was genotyped with 53 co-dominant markers and the linkage mapping was performed on the 1,014 individuals with a low amount of missing data. The introgression size was estimated to be 18.7 cM limited distally by the marker EXCAP_16 and proximally by the marker MACE_b_79 (Figure 2). The resistance locus *Ryd4*^*Hb*^ is flanked by the MACE marker Mace_b_53 and the TC marker TC262452 with a genetic distance of 0.3 cM proximally and 0.5 cM distally, respectively. Moreover, the 200 BC_2_F_5_ plants screened with TC262452 and phenotyped by DAS-ELISA showed a perfect cosegregation of the TC262452 marker data with the resistance: the 100 plants carrying at least one Hb allele at the TC173485 marker had an extinction <0.1 whereas the 100 plants homozygous for the Hv allele had an extinction >0.5.

**Fig. 2.**
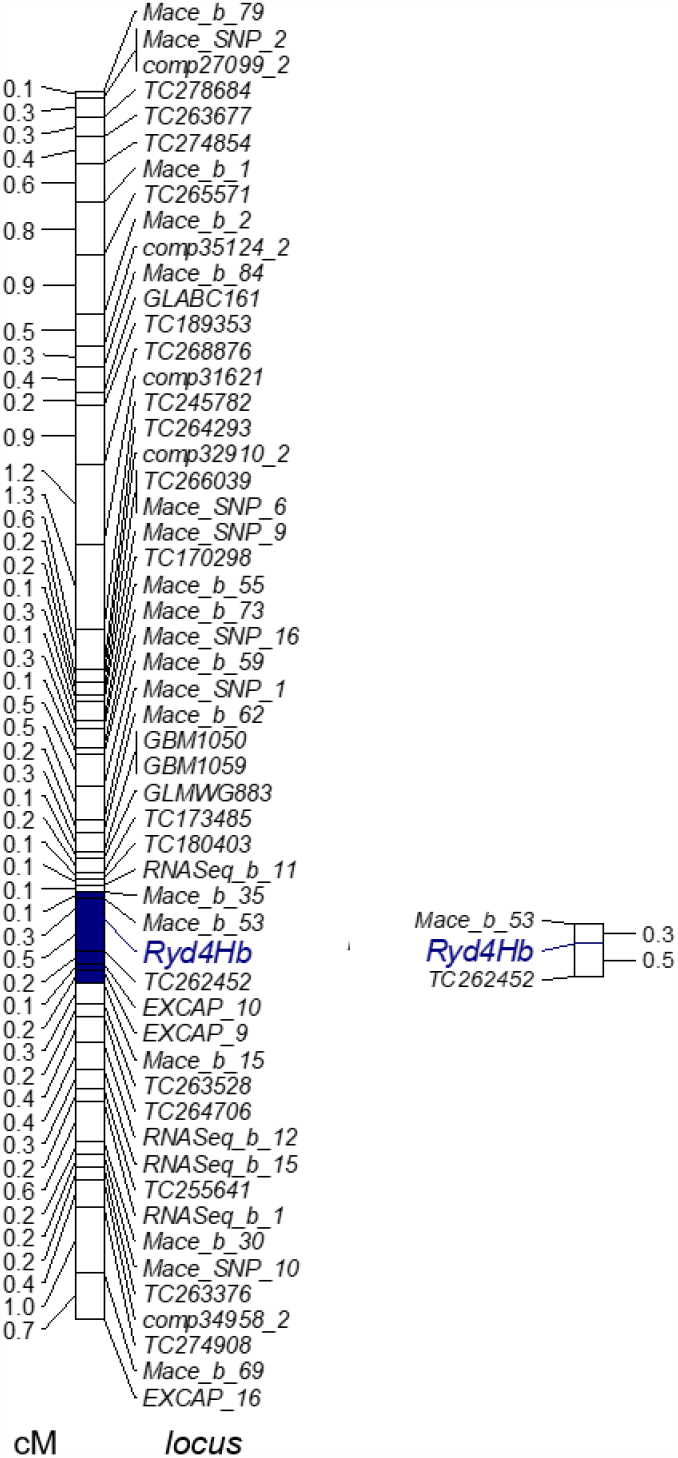
Linkage map of chromosome 3HL carrying *Ryd4*^*Hb*^. The extent of the *Ryd4*^*Hb*^ locus is represented in blue (1), marker bracket around *Ryd4*^*Hb*^ displaying the flanking markers (2)

### Development of an introgression line without sublethality factor

The exploitation of *Ryd4*^*Hb*^ for breeding programs requires the selection of homozygous resistant and vital (by opposition with sublethal) plants. To select recombinants lacking the sublethality factor, 12,133 BC_2_F_6_ plants of LM_Pop were screened with markers TC262452 and Mace_b_53. Among those, 3,103 plants were homozygous for Hb-alleles, 6,020 heterozygous (Hb/Hv), and 3,010 homozygous for *H. vulgare* (Hv) at *Ryd4*^*Hb*^ locus, confirming the 1:2:1 segregation (Chi-Square Goodness of Fit Test: χ^2^= 2.14, *p*=0.34). The 3,103 plants homozygous for Hb-alleles were propagated and the progenies were checked for resistance. A single progeny, denoted as JKI-5215, was both completely resistant and vital. Marker analysis revealed that this population was segregating for Hb- and Hv-alleles distally from *Ryd4*^*Hb*^.

The JKI-5215 population, made of 43 BC_2_F_7_ plants from JKI-5215 progeny, was genotyped with all 16 markers that mapped distally from *Ryd4*^*Hb*^. Markers TC262452 to RNASeq_b_1 were homozygous for Hb-alleles whereas Mace_b_30 and all the remaining markers distally located were segregating in a 1:2:1 fashion (Figure 3). The recombination occurred within the initial 3HL introgression and resulted in a reduced Hb-segment of 3.4 cM and all progenies are completely vital. A BC_2_F_8_ from the family JKI-5215 homozygous for the Hv-segment in the sublethality factor interval was selfed and used as the resistant donor for the FM_Pop2.

**Fig. 3.**
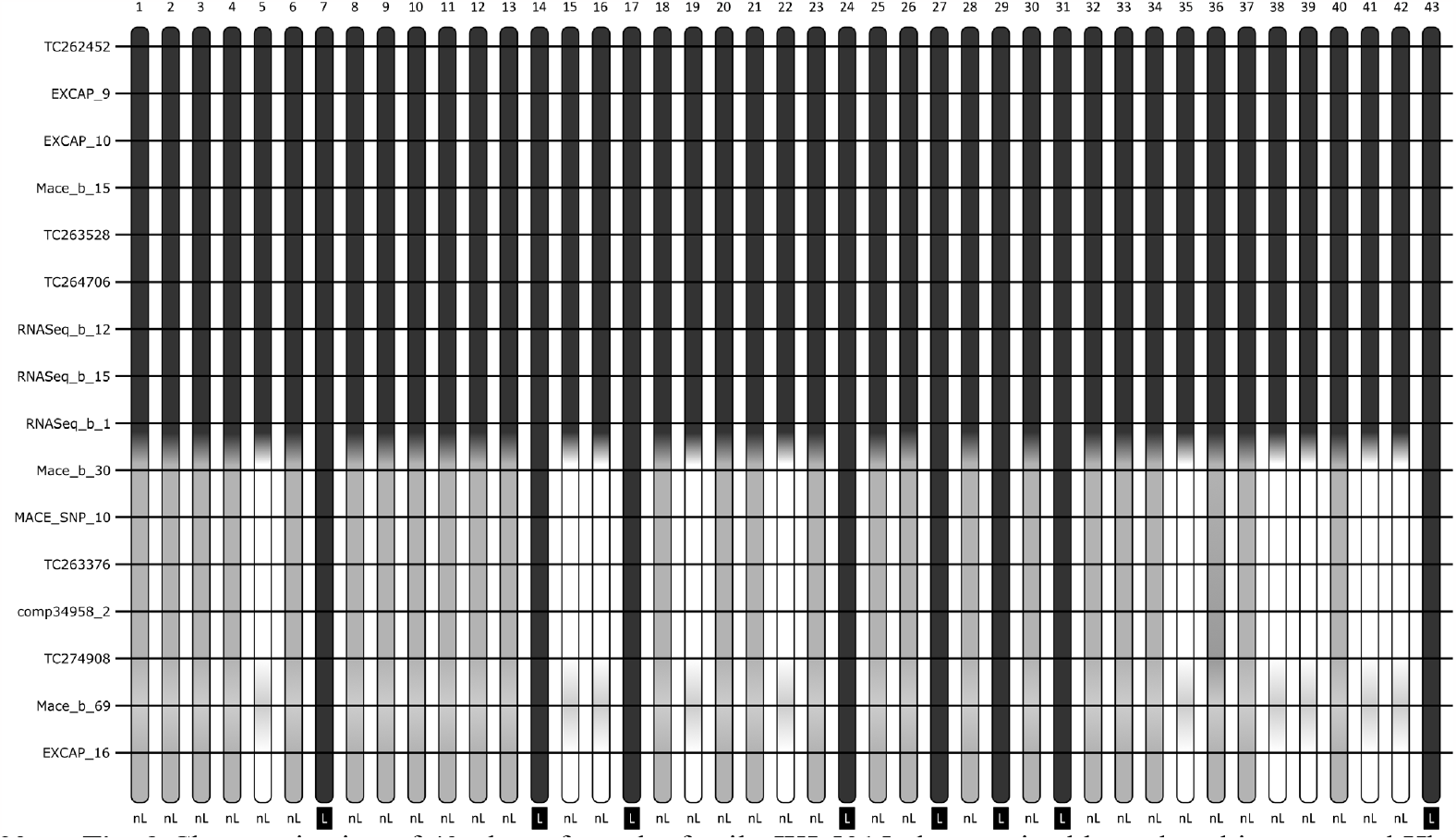
Characterization of 43 plants from the family JKI-5215 characterized by reduced introgressed Hb-fragment without sublethality factor. Each vertical bar represents one offspring from JKI-5215. White, black, and grey fragments represent Hb fragments, Hv fragments, and heterozygous genotypes at the markers indicated on the right side, respectively. The light grey fragments at the dominant marker Mace_b_69 are either Hv or heterozygous genotypes. The phenotype is indicated below the figure as ‘nL’ for vital plants and ‘L’ for sublethal ones.

### High-resolution mapping of *Ryd4*^*Hb*^ in two F_2_ populations

To precisely map *Ryd4*^*Hb*^, 5,589 F_2_ plants from FM_Pop1 and 10,155 F_2_ plants from FM_Pop2 were genotyped with two KASPar markers to identify recombination at the locus. Plants from FM_Pop1 were screened with Ryd4_KASP1 and Ryd4_KASP4, and plants from FM_Pop2 with Ryd4_KASP1 and Ryd4_KASP5. Indeed, Ryd4_KASP4 is located in the sublethality factor interval and was segregating in FM_Pop1, but not in FM_Pop2 where it was fixed Hv in its resistant donor. We identified 46 and 68 recombinant plants in FM_Pop1 and FM_Pop2, corresponding to 0.82% and 0.67% of recombination, respectively. The positions of the SNPs of the markers Ryd4_KASP1, Ryd4_KASP4 and Ryd4_KASP5 were identified in the GBS data from 92 recombinant inbred lines of the cross ‘Barke’ x ‘Morex’ (Mascher et al., 2013). A distance of 15 cM was observed between the markers Ryd4_KASP1 and Ryd4_KASP4, and of 12.6 cM between Ryd4_KASP1 and Ryd4_KASP5, indicating by definition a 15% and 12.6% probability of recombination in these intervals, respectively. The observed rate of recombination at the *Ryd4*^*Hb*^ locus is therefore about 20-times lower than expected for the same genetic interval in a pure intraspecific barley cross.

Recombinants were phenotyped on 15 offsprings and genotyped with the 13 CAPS markers. The resulting interval for *Ryd4*^*Hb*^ was 66.5 kbp-long in MorexV3 genome between the coordinates 592,685,940 and 592,752,329 flanked by CAPS19_2 and CAPS24, describing an interval comprising six recombination events (Figure 4, Supplementary table 7). Four genes are annotated with high confidence on the MorexV3 genome in that interval: HORVU.MOREX.r3.3HG0318400, an S-formylglutathione hydrolase, HORVU.MOREX.r3.3HG0318420, a partial nucleotide-binding and leucine-rich repeat immune receptors (NLR) with a coiled-coil domain (CNL) lacking LRR domain which is likely a pseudogene, HORVU.MOREX.r3.3HG0318450, a complete CNL, and HORVU.MOREX.r3.3HG0318470, an ankyrin-repeat-containing gene. Those genes will later be referred to as *Ryd4_SFGH, Ryd4_pCNL1, Ryd4_CNL2*, and *Ryd4_ANK*, respectively. The genes’ homology revealed that locus *Ryd4*^*Hb*^ is syntenic to the *Triticum monococcum* locus *Sr35*, conferring resistance to wheat stem rust (Saintenac et al., 2013). The candidate genes present a high similarity to one of the *Sr35* candidate genes. In particular, the translation of the complete *Ryd4_CNL2* gene sequence from the MorexV3 reference genome shows 83% identity with the SR35 protein while the respective coding sequence shows 88.7% nucleotide identity. The *Ryd4*^*Hb*^ interval also overlaps almost completely the ones of the *Rph13* leaf rust resistance gene from the *H. vulgare* ssp. s*pontaneum* Hs2986 (Jost et al., 2020) and of the *Jmv2* resistance gene to the *Japanese soil-borne wheat mosaic virus* from the barley cultivar ‘Sukai Golden’ (Okada et al., 2022). *Rph13* is located on chr3H between the coordinates 592,658,337 and 592,786,929 on MovexV3 (128.6 kbp). Comparing the number of recombinants, the size of the interval in which they occurred, and the size of the mapping population for *Rph13* and *Ryd4*^*Hb*^ (4 recombinant in 128.6 kbp out of 719 plants and 6 recombinants plants in 66.5 kbp out of 15,774, respectively), the recombination rate observed in the *Ryd4*^*hb*^ populations is 7.5 times lower than observed in the intraspecific cross used to map *Rph13*.

**Fig. 4.**
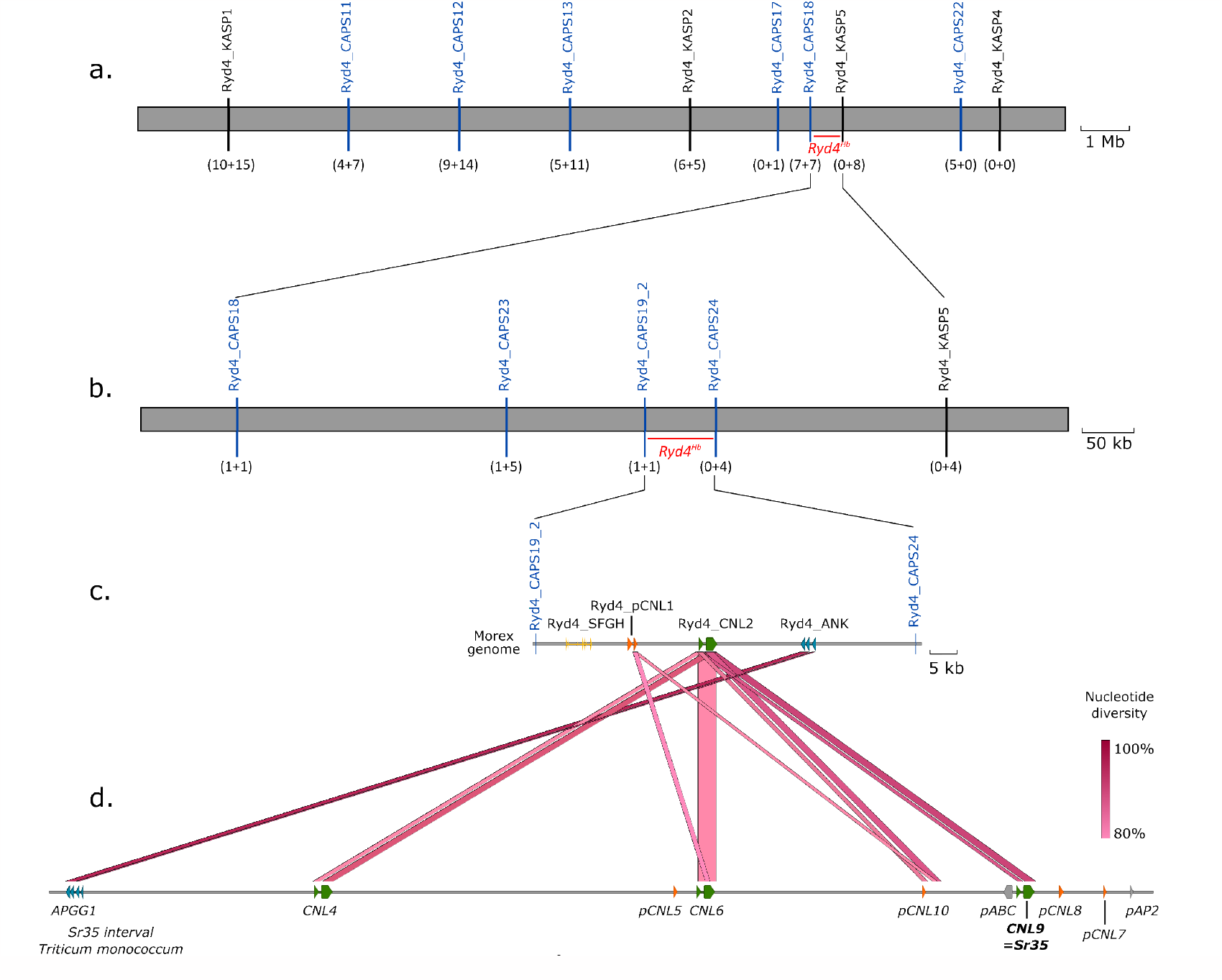
High-resolution mapping of the *Ryd4*^*Hb*^ locus. (a) High-resolution mapping in the 5,589 F_2_ plants from FM_Pop1 and 10,155 plants of FM_Pop2. The numbers in brackets show the sum of individual recombinants between the resistance locus and the corresponding marker in FM_Pop1 and FM_Pop2, respectively. (b) Marker saturation of the 18 recombinant plants at *Ryd4*^*Hb*^ locus. Recombinants from FM_Pop1 that died are excluded. (c) Candidate genes in the 66.5 kbp final *Ryd4*^*Hb*^ interval. (d) Comparison of *H. vulgare* ‘Morex’ and *T. monococcum* DV92 orthologous genes (only fragments with more than 80% nucleotide identity are shown).

### *Ryd4*^*Hb*^ locus diversity in the barley pangenome

The orthologous intervals of the MorexV3 *Ryd4*^*Hb*^ region were retrieved from 19 additional diverse genome assemblies of the barley pangenome (Jayakodi et al., 2020) (supplementary table 8). We then annotated them using a combination of methods: mapping the ‘Morex’ genes, searching for NLR genes with NLR-Annotator (Steuernagel et al., 2020), and confirmation of the absence of additional conserved domains with NCBI conserved domains (S. Lu et al., 2019). The analysis revealed a very large divergence of the *Ryd4*^*Hb*^ interval in the different genotypes (Figure 5, supplementary figure 1). The shortest orthologous interval is the one of MorexV3. The largest is the one of the accession ‘HOR 21599’, 406 kbp-long and containing 10 NLRs, of which five are complete. The interval in the cultivars ‘Akashinriki’ and ‘Du-Li Huang’ is affected by a large inversion of around 600 kbp. The observed diversity between haplotypes is mainly explained by the presence of different repetitive elements and duplications. The degree of divergence in the *H. vulgare* genepool in this interval suggests that an even greater diversity and divergence may be anticipated for the corresponding region in the *H. bulbosum* genome.

**Fig. 5.**
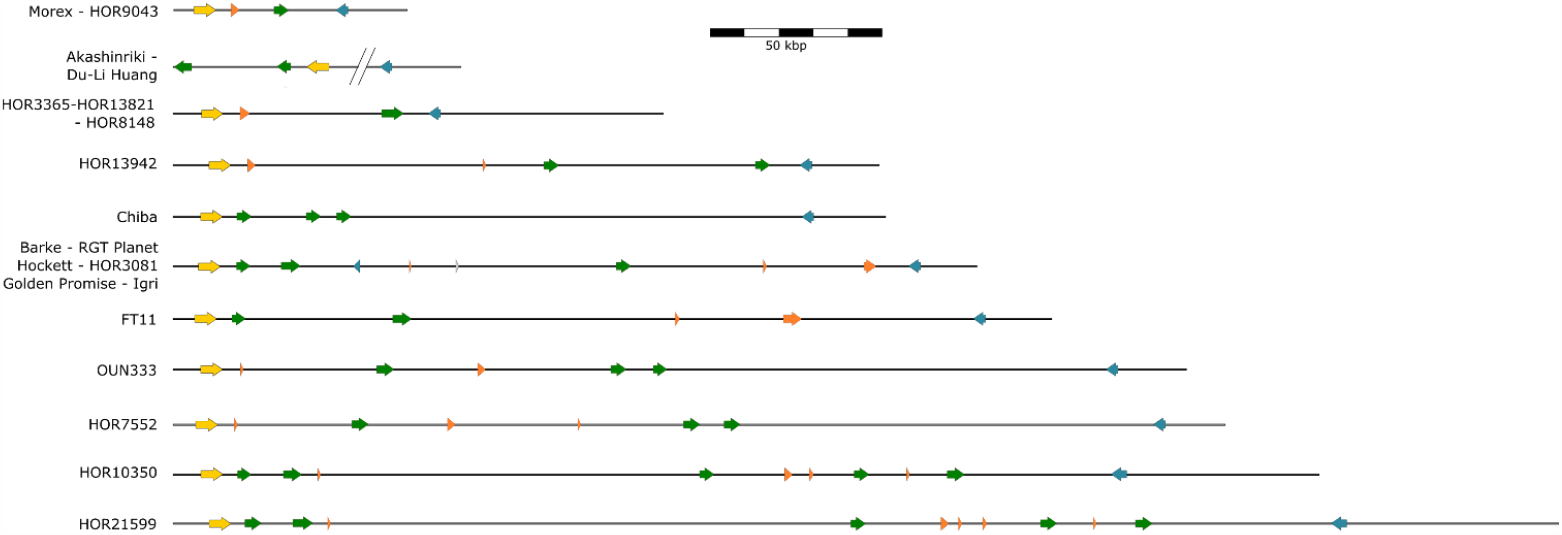
Graphical representation of the haplotype size and gene composition in the barley pangenome. Grey horizontal bars represent the dimension of the interval; forward slashes represent the breakpoint due to the large inversion in Akashinriki and Du-Li Huang. The interval between them is around 400 kbp-long and is not homologous to ‘Morex’ interval. Arrows represent genes; yellow genes are S-formylglutathione hydrolase, orange genes are partial NLRs, green genes are complete NLRs, and blue genes are ankyrin-repeats-containing genes.

### Mapping of the sublethality factor

To better understand why the *H. bulbosum* locus is causing the sublethality, we used recombinants identified in the frame of *Ryd4*^*Hb*^ mapping to pinpoint the responsible factor more precisely. The low-resolution linkage mapping located it distally from the marker *RNASeq_b_1*, which corresponds to position 594,019,595 on chromosome 3H of the MorexV3 genome. To precise its interval, ten PACE markers were designed between Ryd4_KASP18 and Ryd4_KASP22 (supplementary table 5), and used to genotype plants recombining in the interval. Six recombination events were available: the one of JKI-5215 that we mapped using 24 non-recombinant F_2_ plants from lineage EP_16-460 of FM_Pop2, and the ones of five F_2_ plants from FM_Pop1 recombining between Ryd4_KASP5 and Ryd4_C22, of which four vital plant and the sublethal plant KW15_231/340_48 which died before heading (figure 5). The genotyping of 24 non-recombinant FM_Pop2 F_2_ plants confirmed that the JKI-5215 recombination event occurred between Ryd4_KASP5 and Ryd4_CAPS22 (Figure 5), and more precisely between markers Ryd4_leth2 and Ryd4_leth3 (594,034,042 to 594,290,776 bp). The genotyping and phenotyping of thirty-two F_3_ plants from each of the four vital plants recombining between Ryd4_KASP5 and Ryd4_C22 placed the sublethality factor proximally of marker Ryd4_leth7. This was confirmed by the genotyping of the sublethal plant KW15_231/340_48 from FM_Pop1 which was identified as recombinant between Ryd4_leth6 and Ryd4_leth7 (Figure 6). The sublethality factor could therefore be assigned to a 483 kbp-interval between markers Ryd4_leth3 and Ryd4_leth7 (594,290,776-594,773,972 bp on chromosome 3H of MorexV3). This interval is annotated with 15 high-confidence genes described in table 1. Among those genes, one or several could be essential genes for plant development with no orthologs in the corresponding region of the *H. bulbosum* genome.

**Fig. 6.**
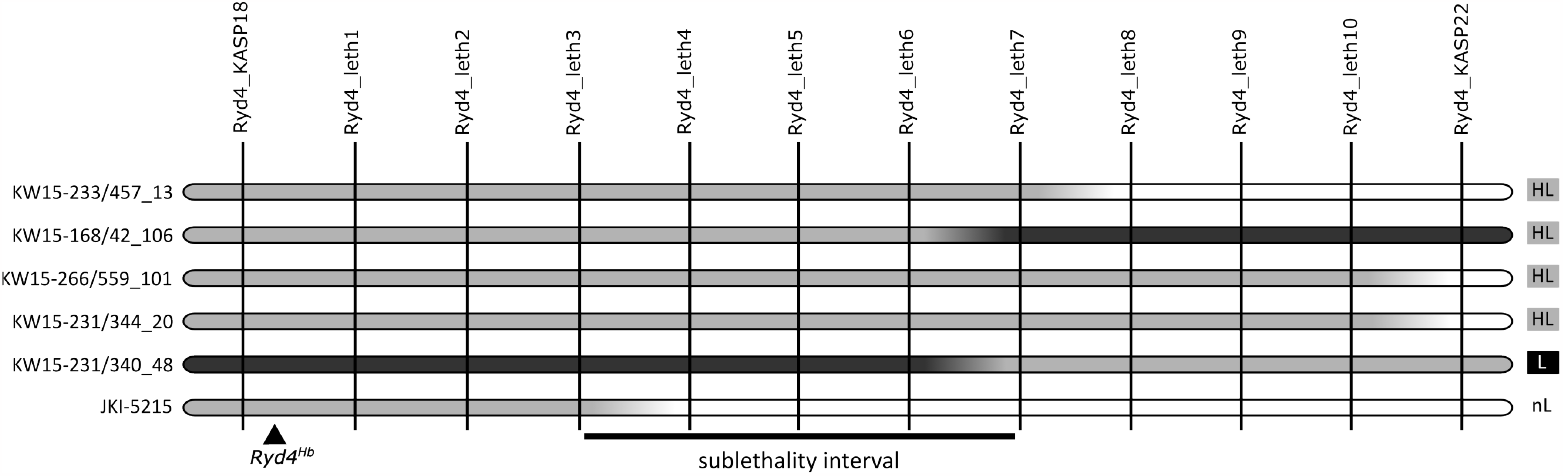
Graphical representation of the genotype of lines recombining in the sublethality interval. Genotypes of KW15-233/457_13, KW15-168/41_106, KW15-266/59_101, and KW15-231/344_20 are inferred from those of 32 of their offsprings. The genotype of JKI-5215 is reconstructed from those of 24 F_2_ plants of FM_Pop2 from the lineage EP_16-460. Markers are depicted as vertical black lines and genotypes as horizontal bars. White, black, and grey segments represent Hv, Hb, and heterozygotes genotypes, respectively. Phenotypes are described on the right as sublethal (L), sublethality segregating in the progeny (HL), and vital (nL). For better readability, marker positions are not to scale.

### *Ryd4*^*Hb*^ does not prevent aphid feeding

Resistance to insect-transmitted viruses can either be a direct resistance to the virus or an indirect one, through resistance to the vector. To test if *Ryd4*^*Hb*^ provides resistance to its aphid vector, we monitored the feeding of 12 to 16 aphids by EPG on five lines: two susceptible F_4_ lines (EP_16-271_3_206_2 and EP_16_460_5_228_2), their two resistant sister lines (EP_16-271_3_206_4 and EP_16_460_5_228_6) and the susceptible barley cultivar ‘Igri’ which was the susceptible parent of LM-Pop and JKI-5215 (Figure 7a). As none of the selected EPG parameters showed a normal distribution according to a Shapiro-Wilk test with a p-value threshold of 0.05, we selected the Kruskal-Wallis test for multiple comparison. No significant differences between the lines were observed for the selected parameters s_Np (*χ*^*2*^=6.78, df=4, p=0.148), s_C (*χ*^*2*^=2.35, df=4, p=0.671), s_F (*χ*^*2*^=4.64, df=4, p=0.327), s_G (*χ*^*2*^=2.96, df=4, p=0.565), s_E1 (*χ*^*2*^=1.35, df=4, p=0.854), s_E2 (*χ*^*2*^=3.16, df=4, p=0.534) and s_sE2 (*χ*^*2*^=3.52, df=4, p=0.474) (Figure 7b, supplementary table 9). The most divergent line was Igri, with an increased median duration for s_Np and decreased median durations for s_E2 and s_sE2, probably due to differences in the genetic background with the other lines.

**Fig. 7.**
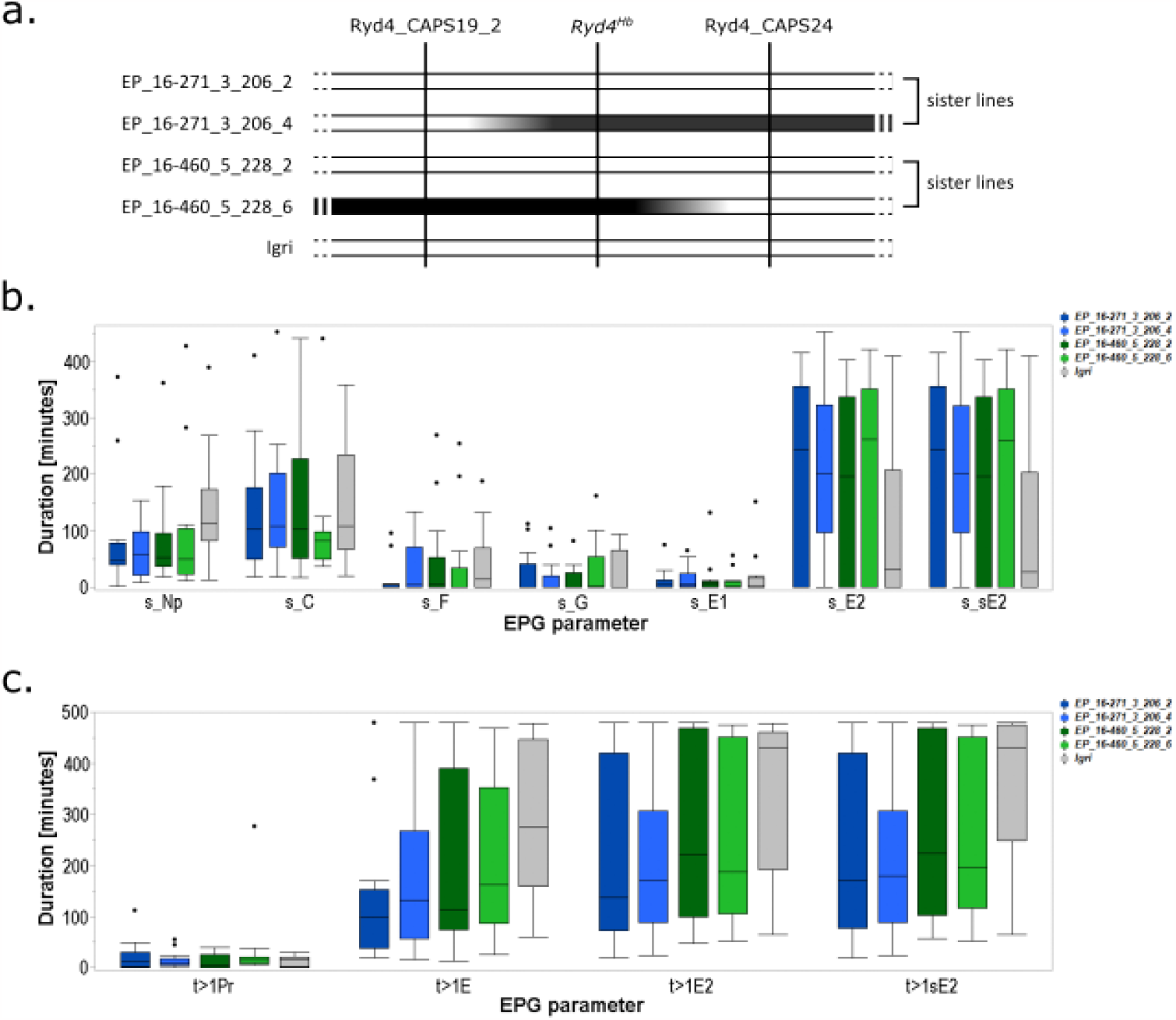
Aphid feeding behaviour on resistant and susceptible lines **a**. Graphical representation of the lines used to test the aphid feeding behaviour on resistant and susceptible lines. Loci are depicted as vertical black lines and genotypes as horizontal bars. White and black, segments represent Hv and Hb genotypes, respectively. **b**. Duration of the different feeding events on the five lines summed up by type. s_Np: total duration of all non-probing events. s_C: total duration of pathway phase. s_F: total duration of penetration problems. s_G: total duration of xylem drinking. s_E1: total duration of secretion of watery saliva. s_E2: total duration of phloem sap ingestion. s_sE2: total duration of sustained phloem sap ingestion. **c**. Duration … Lines in the box plots indicate the median, and whiskers show the upper and lower 1,5xIQR (interquartile range) with dots indicating ouliers.

In addition to these general parameters, parameters associated with epidermis (t>1Pr) and mesophyll located (t>1E) but also sieve element located (t>1E2, t>1sE2) defense responses were inspected. Igri showed the\ longest median durations for reaching the sieve elements (t>1E) and to reach ingestion (t>1E2 and t>1sE2). However, none of the parameters t>1Pr (*χ*^*2*^=3.83, df=4, p=0.43), t>1E (*χ*^*2*^=6.32, df=4, p=0.176), t>1E2 (*χ*^*2*^=3.43, df=4, p=0.489) and t>sE2 (*χ*^*2*^=5.42, df=4, p=0.247) differs significantly between the tested lines (Figure 7c, supplementary table 9).

## Discussion

BYDV is a major threat to barley cultivation that is expected to increase in the following years, as autumns become longer and warmer in Northern Europe (Roos et al., 2011; Trebicki, 2020), thus breeding for BYDV resistance will become an even higher priority. So far only partial resistance has been discovered in the *H. vulgare* primary genepool (Baltenberger et al., 1987; Collins et al., 1996; Lüpken et al., 2014; Niks et al., 2004; Schaller et al., 1964; Suneson, 1955). Here, we report the high-resolution mapping of *Ryd4*^*Hb*^, the first resistance gene to BYDV in barley, originating from the wild relative and secondary genepool species *H. bulbosum*. We mapped the gene in a 66.5 kbp-interval of the MorexV3 barley reference genome despite the linkage of the resistance locus to a sublethality factor and the reduced recombination between *H. vulgare* and *H. bulbosum* genomes.

At the *Ryd4*^*Hb*^ locus, four genes are annotated with high confidence on the MorexV3 genome, including two genes from the CNL family, one pseudogene, and one likely to be functional. CNL genes are part of the larger NLR family which are the most common class of resistance genes to biotic stress. They are coding for intracellular proteins that form complexes (Wang et al., 2019) recognizing, directly or indirectly, pathogen effector molecules and inducing local cell death in response. More than 30 NLR genes conferring resistance to viruses have been cloned so far (Boualem et al., 2016; Sett et al., 2022) and more are candidates. An *H. bulbosum* homolog of *Ryd4_pCNL1* or *Ryd4_CNL2* is therefore a very likely candidate for *Ryd4*^*Hb*^. The *Ryd4*^*Hb*^ locus is orthologous to the *Sr35* resistance locus from the wheat wild relative *Triticum monococcum* (Saintenac et al., 2013). *Sr35* is coding for a CNL protein that shares 83% identity with the translated *H. vulgare* sequence of *Ryd4_CNL2* and provides resistance to the fungal pathogen *Puccinia graminis* f. sp. *tritici* causing wheat stem rust. This interval was also identified as candidate for the *Rph13* resistance gene to leaf rust in the *H. vulgare* spp. *spontaneum* accession ‘PI 531849’ (Jost et al., 2020), and, overlaps with the large interval of the *Jmv2* resistance gene to the *Japanese soil-borne wheat mosaic virus* from the barley cultivar ‘Sukai Golden’ (Okada et al., 2022). Interestingly, the best homolog of *Sr35* in rice is LOC_Os11g43700, which was identified as a resistance gene to the *Rice yellow mottle virus* in the African rice species *Oryza glaberrima* (Pidon et al., 2017; Bonnamy et al., 2023). It is not rare that closely related NLRs provide resistance to different classes of pathogens. A good example is the potato NLRs genes GPA2 and RX1 which provide resistance against the nematode *Globodera pallida* and potato virus X, respectively, and share 88% of their amino acid sequence (Van Der Vossen et al., 2000). The comparison of the *Ryd4*^*Hb*^ interval in the barley pangenome (Jayakodi et al., 2020) demonstrated a very large diversity at this locus, including NLR duplications. NLRs genes are indeed frequently under diversifying selection and tend to evolve and duplicate by interallelic recombination between orthologs and by unequal crossing-over (Baggs et al., 2017; Chen et al., 2010; Ding et al., 2007; Guo et al., 2011; Li et al., 2010; Michelmore & Meyers, 1998; Zhou et al., 2004). The *Sr35/Ryd4*^*Hb*^ locus is one of those very diverse and dynamic loci that could be described as R gene factories. Together with the homology with other resistance loci, this locus’ NLR diversity pleads for *Ryd4*^*Hb*^ to be a CNL. However, in addition to the CNL genes, an *H. bulbosum* ortholog of the *Ryd4_ANK* could also be a good candidate. Indeed, the structure of the encoded protein is close to the one of *Arabidopsis* ACCELERATED CELL DEATH 6 (ACD6) protein. ACD6 confers enhanced resistance to bacterial pathogens, including *Pseudomonas syringae*, by increasing the level of salicylic acid and inducing spontaneous cell death (Rate et al., 1999; Dong, 2004; H. Lu et al., 2003, 2005). We also cannot exclude that *Ryd4*^*Hb*^ resistance is due to presence/absence variation of a gene in the primary and the secondary genepool of barley, thus the resistance gene from *H. bulbosum* may have no ortholog in the *H. vulgare* interval. Cloning of *Ryd4*^*Hb*^ would therefore most likely require a *de novo* genome assembly of the *Ryd4*^*Hb*^ interval in a resistant genotype (introgression line of *H. vulgare* or resistance donor genotype of *H*.*bulbosum*).

Resistance to insect-transmitted viruses, like the one provided by *Ryd4*^*Hb*^, can either be a direct resistance to the virus or a resistance to the vector, which would in effect prevent infection and therefore provide indirect virus resistance. The melon NLR *Vat* resistance gene is the model of this indirect resistance. VAT provides resistance to *Aphis gossypii* and to all the viruses it transmits tested so far, including the *Cucumber mosaic virus* (Boissot et al., 2016). It recognizes an effector from *A. gossypii* and triggers the hypersensitive response, stopping at the same time any viral infection that may have occurred. BYDV cannot be inoculated to barley mechanically, so only resistance to the aphid vector was tested. A previous study showed that *R. padi* aphids were feeding less and having a shorter salivation time on the *Ryd4*^*Hb*^ *H. bulbosum* resistance donor A17 compared to the BYDV-susceptible *H. bulbosum* line A21, suggesting that this could be the reason for A17 BYDV resistance (Schliephake et al., 2013). However, our study showed no differences in aphid feeding patterns on closely related resistant and susceptible lines, accompanied by an absence of BYDV infection in the resistant lines, suggesting that the preliminary observation on the *H. bulbosum* donor was probably due to A17 genetic background rather than *Ryd4*^*Hb*^. *Ryd4*^*Hb*^, therefore, provides direct resistance to BYDV.

*Ryd4*^*Hb*^ is a prime example of the importance of crop wild relatives serving as genetic resources and gene donors in breeding schemes to achieve efficient and durable disease resistance. The advantage of using a crop wild relative in prebreeding schemes as a unique source of resistance, however, comes at a cost: reduced frequency of recombination and/or hybrid incompatibility leading to fertility or lethality problems. In the case of the *Ryd4*^*Hb*^ locus, recombination is 7.5 times less than in the intraspecific barley cross used to map *Rph13*. To fine map the gene despite this handicap, we screened very large mapping populations with high throughput genotyping technologies. At the *Ryd4*^*Hb*^ locus, this negative linkage drag was strongly materialized by a sublethality factor characterized by the reduced growth and early death of introgression lines carrying the Hb allele at homozygous state at this locus. By screening a large number of plants for recombination, we managed to break the linkage, producing a resistance donor without the sublethality factor, that could be included in breeding schemes. We mapped the sublethality factor to a 600 kbp interval on MorexV3 genome. The observed phenotype suggested that sublethal plants are possibly lacking an essential gene, or a few genes, for development, therefore that one of the genes of the *H. vulgare* interval has no ortholog in the donor *H. bulbosum* genome. Among the 15 genes annotated with high confidence in the interval, B3 domain-containing proteins are part of a large transcription factor superfamily whose members are playing key roles in various stages of plant development, from embryogenesis to seed maturation (Swaminathan et al., 2008). F-box containing proteins are central part of the ubiquitin–26S proteasome system and are thus key for different processes like phytohormone signaling, plant development, cell cycle, or self-incompatibility (Stefanowicz et al., 2015). RecA proteins are maintaining DNA integrity during meiosis by initiating double-strand break repair (Emmenecker et al., 2023). Annexins are widely involved in regulating plant processes, from growth and development to responses to stresses (Wu et al., 2022). One of the corresponding genes in the MorexV3 interval could be missing in the A17 haplotype and explain the observed phenotype.

The results of this study would be helpful to breed barley varieties with an effective resistance to BYDV. We identified recombinants with a strongly reduced *H. bulbosum* fragment that can be used in breeding schemes, removing almost completely the negative linkage drag. The markers closely linked to the resistance can then be used in marker-assisted and genomic selection, postponing the tedious resistance evaluation to the last breeding step. Knowing that *Ryd4*^*Hb*^ is a direct resistance gene to BYDV would also make it possible to establish the best strategy to avoid resistance breaking. Such a strategy could be pyramiding it with partial resistance or tolerance sources like *Ryd2* and *Ryd3*. Such resistant varieties would make a major contribution to sustainable barley cultivation.

## Declarations

### Availability of data and material

The Exome capture sequencing datasets generated and/or analyzed in this study are deposited at EMBL-ENA under the project IDs PRJEB7909 and PRJEB65283.

### Authors’ contributions

Brigitte Ruge-Wehling, Nils Stein, Neele Wedler, and Viktor Korzun conceived the project and acquired the funding. Brigitte Ruge-Wehling, Neele Wedler, Klaus Oldach, Anja Maasberg-Prelle, and Viktor Korzun designed and built the mapping populations. Brigitte Ruge-Wehling and Kristin Fischer performed the initial linkage mapping and broke the linkage between the resistance and the sublethality loci. Brigitte Ruge-Wehling and Antje Habekuß performed the phenotyping experiments. Neele Wedler did the exome capture experiment. Torsten Will performed and analyzed the aphid feeding experiments. Hélène Pidon performed the high-resolution mapping and the pangenome analysis and drafted the manuscript. Nils Stein supervised the project. All authors provided critical feedback and helped shape the manuscript.

## Acknowledgments

We gratefully acknowledge the excellent technical support by Manuela Kretschmann in DNA extraction and KASP genotyping, Dörte Grau in BaMMV resistance phenotyping, and Evelyn Betke, Katharina Stein and Kerstin Welzel in aphid rearing. We thank Anne Fiebig for raw data submission and Kristin Kutter for marker analysis.

**Supplementary figure 1** Comparison of *Ryd4*^*Hb*^ in *H. vulgare* pangenome modify from Easyfig 2.2.5 output. Only fragments with more than 80% nucleotide identity are shown.

