## Supplementary figures and images for "High-resolution mapping of *Ryd4^Hb^*, a major resistance gene to *Barley yellow dwarf virus* from *Hordeum bulbosum*"

### Supplemental figure 1

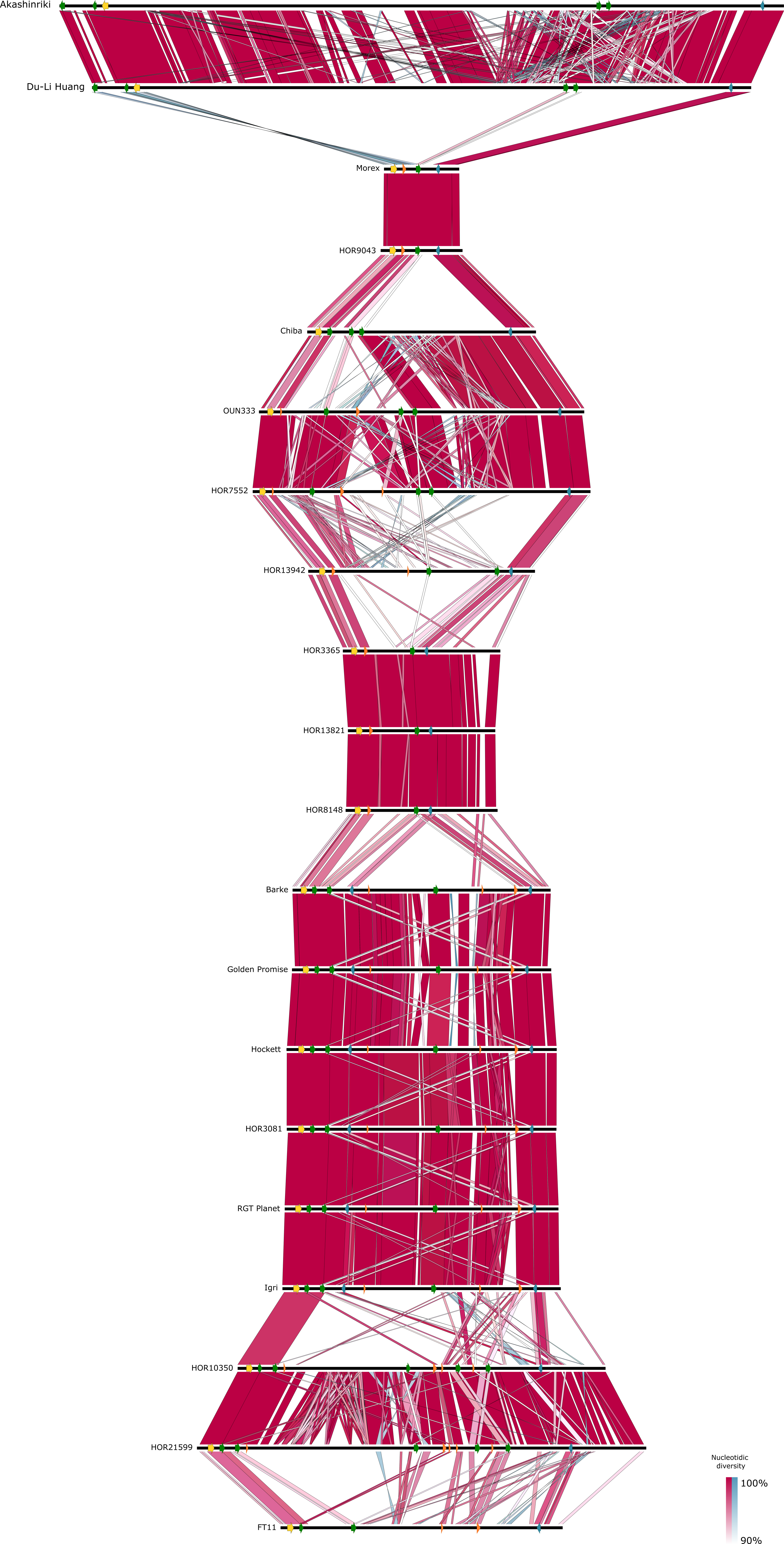
